# Efficient Data Collection for Establishing Practical Identifiability via Active Learning

**DOI:** 10.1101/2025.07.28.667128

**Authors:** Xiaolu Liu, Linda Wanika, Michael J. Chappell, Juergen Branke

## Abstract

Practical identifiability analysis (PIA) plays a crucial role in model development by determining whether available data are sufficient to yield reliable parameter estimates. In bioengineering applications, identifying the minimal experimental design that ensures parameter identifiability is essential in order to reduce cost, time, and resource consumption. In this paper, we introduce E-ALPIPE, a sequential active learning algorithm that recommends new data collection points most likely to establish practical identifiability given the current data, mathematical model and noise assumptions. We empirically evaluate E-ALPIPE against both a benchmark algorithm from the literature and random sampling over three synthetic experiments. Our results show that E-ALPIPE requires up to 50% fewer observations on average to achieve practical identifiability, compared to the strongest competitor, while producing comparable or narrower confidence intervals and more accurate point estimates of system dynamics.

**Author summary:** Scientists and engineers often use computer models to understand complex biological systems, such as how cells grow or how chemicals behave in a reactor. To make these models useful, they need to determine the correct values for various model parameters, based on data from experiments. However, experiments are usually expensive, time-consuming, and often produce noisy data. If the available data is insufficient, our confidence in the fitted model parameters may be low and thus the model’s predictions might be unreliable. This is known as a problem of practical identifiability.

This paper introduces a new method called E-ALPIPE, which helps scientists decide which experiment is likely to produce the most useful information for improving their model. Using E-ALPIPE they can determine the model parameter values more efficiently, saving time and resources.

We tested E-ALPIPE on several example problems and found that it could reach the same level of reliability using up to 50% fewer experiments than standard methods. It also produced more precise and accurate results. Overall, E-ALPIPE is a smarter, more efficient way to design experiments for tuning the parameters of computer models.

## Introduction

Bioengineering models have facilitated significant advancements in biological understanding and medical insights [1–3]. The use of hybrid data-driven and mechanistic models has increased in popularity over recent years [4]. This pairing enables model parameters to be estimated from observed experimental outputs, and allows modellers to check whether the model simulations are closely aligned with the actual phenomena characterised by the model.

A prerequisite for reliable parameter estimation is identifiability. Structural identifiability analysis (SIA) determines whether unique parameter values are theoretically recoverable from noise-free, continuously sampled data. If a model is structurally unidentifiable, infinitely many parameter values can reproduce the same model output. Practical identifiability analysis (PIA) extends the concept to finite, noisy datasets, assessing whether parameters can be estimated with acceptable uncertainty under realistic experimental conditions [5, 6]. Consequently, a model is practically unidentifiable when many parameter sets yield statistically indistinguishable outputs. A common approach to rendering a practically unidentifiable model identifiable is through the collection of additional data [7].

While experimental data inform models, well specified models can guide data acquisition as well. Yet, as [6] state, “… there remains a persistent lack of generalisable methods to optimise sample placement and experimental design.”

Several prior studies have incorporated PIA into experimental design workflows [7–11], typically combining profile likelihoods with sensitivity analyses to guide sampling design. However, these approaches often require extensive model evaluations, and may assume access to unrealistic levels of data. For example, [9] assumes access to an existing complete experimental dataset to identify minimally sufficient experiments. While insightful, it is not feasible as a general experimental design strategy in practice. More generally, computing profile likelihoods for complex models remains a significant challenge [11]. Simulating data across all time points and evaluating likelihoods over a wide parameter space is computationally expensive. A state-of-the-art algorithm proposed in [10]—and applied in [12, 13]—also aims at optimal experimental design, but it relies on a thorough profile-likelihood analysis over the entire parameter space in each iteration. This algorithm is included as a benchmark in our experimental comparisons.

To tackle the problem of generalisable and low-cost practical identifiability-driven design, we propose the Efficient Active Learning Practical Identifiability Parameter Estimation algorithm (E-ALPIPE), an active-learning algorithm that iteratively selects the next experiment with the goal of achieving practical identifiability using a minimal number of measurements. E-ALPIPE allows for measurement noise and can be applied to both linear and non-linear ordinary differential equation models. Its performance is compared against the benchmark algorithm proposed in [10] as well as random sampling. We evaluate the algorithm on three synthetic case studies: one drawn from the benchmark study of [12] and two based on a bioreactor model, one of those with two output dimensions.

The structure of this paper is as follows: Section 1 reviews the theoretical foundations. Section 2 details our E-ALPIPE algorithm, while the benchmark algorithm is described in Section 3. Empirical results are presented in Section 4. Finally, Section 5 discusses limitations, contributions, and directions for future research.

## 1 Background

This section provides a brief introduction to the principles of uncertainty sampling in active learning as well as practical identifiability analysis using the profile likelihood approach.

### 1.1 Uncertainty sampling within active learning

Active Learning is an influential paradigm in both supervised and semi-supervised machine learning, and is especially effective in optimising model performance under a restricted budget [14]. It is notably beneficial in scenarios with high costs for acquiring labelled data [15]. Its goal is to maximise model performance under a limited labelling budget by selecting only the most informative unlabelled instances for annotation. The criterion for what constitutes informative data varies across algorithms, but universally aims to enhance model accuracy with fewer labelled instances, thereby minimising the associated costs [16].

In this work, we focus on sequential active learning, which iteratively chooses a single new sample based on the current model’s uncertainty [14]. Among the various data selection criteria, Uncertainty Sampling is the most widely studied: at each step, the algorithm queries the point whose predicted label is least certain under the model, then updates the model with that newly labelled data [14].

Common measures of prediction uncertainty include variance, least confidence and entropy [14]. However, the detailed interpretation of uncertainty varies depending on the specific requirements of different research approaches [17]. In Section 2 we propose a novel metric that can take into account varying noise levels and multiple outputs.

### 1.2 Practical identifiability analysis – profile likelihood approach

Practical identifiability analysis determines whether model parameters can be reliably estimated from the available experimental data. Various methods exist for assessing practical identifiability, including Markov Chain Monte Carlo (MCMC) approach, the Fisher matrix approach, and profile likelihood analysis [18]. The profile likelihood approach is a common method of practical identifiability analysis employed in biological and bioengineering models [7, 11]. We adopt this method in this paper.

This approach systematically examines how the likelihood function changes as one parameter varies while optimising all other parameters to best fit the data. By analysing the shape and width of the resulting profile likelihood curves, one can evaluate the precision and uniqueness of parameter estimates.

The likelihood of a parameter can be interpreted as the probability of generating the observed output given that parameter, i.e.,

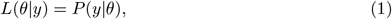

where *θ* refers to a parameter vector, *y* is the observed output, *L* is the likelihood, and *P* is the associated probability.

In many hybrid modelling and data-driven approaches, likelihood serves as a basis for parameter estimation. Modifications in parameter estimates can change the overall likelihood, which is the basis of the profile likelihood approach. Briefly, each unknown parameter has a Maximum Likelihood Estimator (MLE) and an associated likelihood value.

To compute the profile likelihood for a parameter *θ*_*i*_ of interest, we discretise *θ*_*i*_ while re-optimising the remaining nuisance parameters *θ*_*ν*_ to maximise the likelihood for each fixed value of *θ*_*i*_. If *θ*_*i*_ is practically identifiable, we expect a significant change in the likelihood when *θ*_*i*_ is varied. Eq. 2 describes how the 95% profile likelihood confidence interval is computed, where *θ*_MLE_ is the MLE of parameter *θ*_*i*_, *nLL*(*θ*_MLE_) is the negative log likelihood of the MLE, and *χ*^2^(0.95, 1) is the 95th percentile of the chi-squared distribution with one degree of freedom, as the calculated confidence intervals are point-wise based for each parameter [7]:

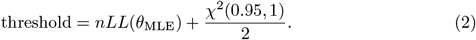

In a typical practical identifiability analysis, one discretises *θ*_*i*_, records the respective likelihoods, and then applies Eq. 2. If the profile likelihood values exceed the threshold on both sides of the MLE, we conclude that the parameter is practically identifiable [7]. The values of *θ*_*i*_ at which the likelihood crosses the threshold on each side of the MLE define the 95% confidence interval.

Since the discretised values of *θ*_*i*_ play an important role in the generated likelihood values and identifiability conclusions, the discretisation strategies are important. Standard tools use fixed or adaptive grids (e.g. 100 points per side), which can be costly [19–21]. To address this, we introduce a targeted bisection-based boundary search (Section 2.4) that reduces computational overheads while preserving accuracy in practical identifiability assessments.

For a more in-depth introduction to practical identifiability analysis, we refer to [5, 7].

## 2 The Efficient Active Learning Practical Identifiability Parameter Estimation (E-ALPIPE) Algorithm

As other active learning approaches to experimental design, our approach works with a set of candidate models and looks at the variability or disagreement between their predictions. So we first discuss these two components, the selection of candidate models and the disagreement metric, before we explain the algorithm in detail. We also propose an efficient search strategy to accelerate practical identifiability assessment.

### 2.1 Candidate models

E-ALPIPE considers a set of candidate models in each iteration which are derived from a simplified profile likelihood analysis.

For a candidate model with a parameter vector *θ*, and assuming independence across existing observations *D* = {(*y*_1_, *x*_1_), (*y*_2_, *x*_2_), …, (*y*_*n*_, *x*_*n*_)}, the joint likelihood of any candidate *f* (*x*|*θ*) is defined by

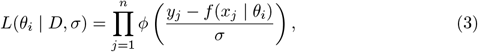

where *ϕ* and Φ denote the standard normal PDF and CDF, respectively. Since observations are assumed to follow a normal distribution truncated below at zero, with known standard deviation *σ*, this becomes

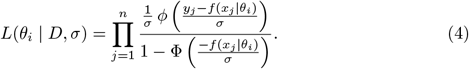

The first candidate model E-ALPIPE considers is then the maximum likelihood model, which is obtained by maximising Eq. 4 over *θ*. The other candidate models are the boundary models from profile likelihood analysis, which are obtained by fixing one parameter at one of its boundary values, and maximising the likelihood over all the other parameters. In total, this generates *m* = 2*N* + 1 candidate models given *N* parameters.

### 2.2 Disagreement among candidate models

As discussed above, the disagreement among possible candidate models is a good indicator for where a sample may be informative. In our setting, in each iteration, we consider *m* candidate models *f* (*x* | *θ*_1_), …, *f* (*x* | *θ*_*m*_), each defined by a distinct parameter vector *θ*_*i*_. However, not all candidate models are equally likely, and thus we weigh their influence based on likelihood, which quantifies how well a model explains observed data.

By normalising the likelihood values across all candidates, we obtain relative weights

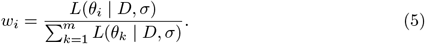

These weights are updated at each iteration and guide the selection of the next experimental design.

With these weights, we can compute, for each candidate location *x*, a weighted disagreement between the boundary models and the MLE model as:

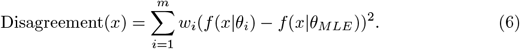

To account for differences in measurement uncertainty across observations, we follow the general principle of the signal-to-noise ratio (SNR) to normalise the information gain. Specifically, at each iteration, we first compute the most likely system trajectory using the current MLE estimate. For each observable, we then estimate the corresponding noise variance based on the current MLE and the known noise model (see Appendix A for details). The weighted disagreement is then normalised by dividing it by the predicted noise variance at each sampling location. Finally, for each candidate location, these normalised disagreements across *M* output dimensions are aggregated to obtain an overall acquisition score, defined by:

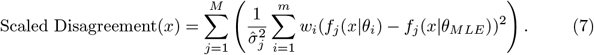

### 2.3 Algorithm

Given the functional form of both the ordinary differential equation (ODE) and noise models, the algorithm begins with the initialisation, and then alternates between two phases:

1. *The estimation phase*: Identifies the Maximum Likelihood Estimation (MLE) model and the boundary models as explained in Section 2.1.
2. *The exploration phase*: Estimate the noise level at each possible sampling location and determine the next data collection point by selecting the point with the highest weighted relative disagreement among the predicted outputs from all the models generated during the estimation phase.

We iterate between the estimation phase and the exploration phase, until practical identifiability is established. The algorithm is summarised in Algorithm 1, with the details as follows:

#### Initialisation (Line 1-2)

We start with a set of random observations, noise model and the analytical form of the function.

#### Estimation (Line 3-10)

With existing observations, we calculate an approximate MLE, and construct a set of boundary models. Each boundary model is found by fixing one parameter at its corresponding boundary value, and maximising the likelihood by optimising the other parameters using the commonly employed local optimisation algorithm L-BFGS with multi-start [22].

#### Decision on practical identifiability (Line 11-17)

From the identified MLE parameters, we can construct the threshold for deciding practical identifiability based on Eq. 2. If all parameters are practically identifiable, and the last iteration was practically identifiable as well, we search for its profile likelihood confidence interval using a bisection search detailed in Section 2.4.

#### Searching for the next experiment (Line 18-25)

We calculate and normalise the likelihood of all candidate models, obtaining a relative probability of each model being the true underlying model (see Section 2.2). Based on these weights, we compute the disagreement score for each location and select the one with the highest overall disagreement.

#### Data collection (Line 26)

We collect new data at the selected location (for all observed outputs) and add these to the dataset to inform the subsequent iterations.

#### Termination (Line 16 and 27)

We iterate until practical identifiability has been established for two consecutive iterations to ensure stability of the results, or until we reach the maximum number of iterations (in the event that practical identifiability has not been reached).

##### Algorithm 1

Efficient Active Learning for Practical Identifiability Parameter Estimation

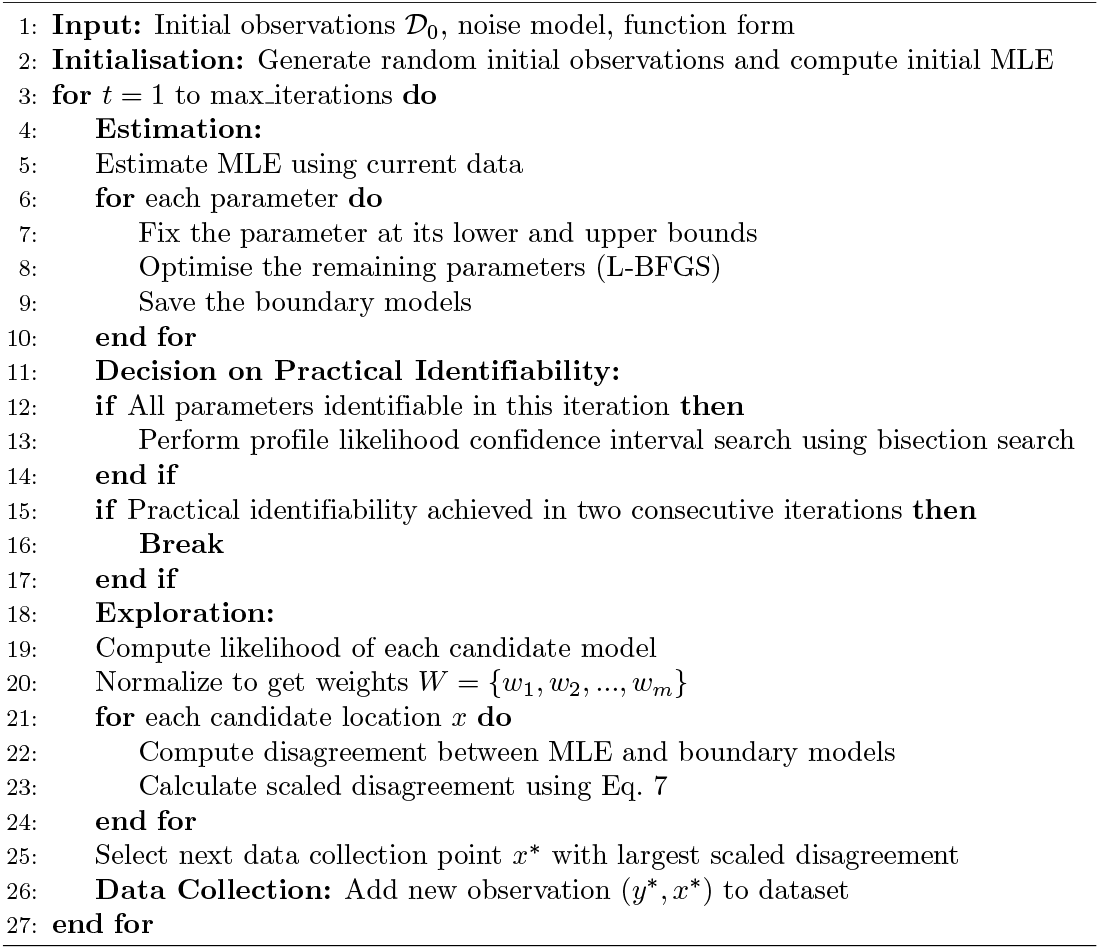

### 2.4 Boundary search for profile likelihood confidence interval

Common practical identifiability analysis tools for bioengineering models are the `data2dynamics` modelling environment in `MATLAB` [19], the `LikelihoodProfiler` package in `JULIA` [21], and the `pyPESTO` package in `PYTHON` [23]. All of these tools and packages use the profile likelihood approach. In `data2dynamics`, for example, default profile likelihood computation explores up to 100 discretised values on either side of the MLE [19]. However, in some instances the default value of 100 steps is not sufficient to accurately identify the profile likelihoos confidence intervals. To enhance the speed and accuracy of computing the profile likelihood confidence intervals, and assuming the profile likelihood is unimodal, we propose to use bisection search [24]. The algorithm works as follows:

- **Initialise:** With the MLE for the given data, we calculate the threshold for the practical identifiability check. Eq. 2 shows the threshold value is only dependent on the confidence level and the MLE, and therefore this threshold can be shared for all parameter settings for which we are checking practical identifiability. We define the initial boundaries to be the user-defined parameter boundaries, and the error tolerance to be a user-defined value *ϵ*_*i*_ for each parameter.
- **Practical identifiability check:** For each parameter, we calculate the profile likelihood *pl*_*min*_ and *pl*_*max*_ at its boundaries. If either *pl*_*min*_ ≤ *threshold* or *pl*_*max*_ ≤ *threshold*, we conclude that this parameter is not (yet) practically identifiable.
- **Profile likelihood confidence interval calculation:** Two bisection searches are performed: one between the parameter’s lower bound and the MLE, and the other between the MLE and the upper bound. These searches iteratively refine the interval and return the estimated lower and upper bounds that define the confidence interval for the parameter under evaluation [25]. If the profile likelihood is continuous, the bisection search will locate a boundary where the likelihood crosses the confidence threshold. In the case of a monotonic profile likelihood, the algorithm guarantees the smallest interval satisfying the confidence criterion.

In practice, we found that it only takes 2-13 iterations to find the intersection point on each side of the parameter estimate, i.e., much fewer than traditional methods.

## 3 Benchmark algorithm

The benchmark algorithm that we compare to has been proposed in [10] and, to the best of our knowledge, is the only other algorithm that explicitly addresses experimental design for improving practical identifiability using profile likelihood. The method involves the use of the profile likelihood trajectories of the parameters to guide experimental planning. Specifically, it determines model outputs from a number of models along their profile likelihood paths, i.e., when each parameter is fixed at a range of values with the other parameters re-optimised. Then, it suggests to take an additional measurement where the predicted outputs of these models vary the most, as they are more likely to reduce uncertainty and tighten the confidence intervals associated with practically non-identifiable parameters [10]. [13] and [12] demonstrate that the approach is practical in real world experimental settings. In their framework, for each candidate experiment, the model is simulated across all acceptable parameter settings derived from the profile likelihood trajectories, and the experiment yielding the widest predicted output range is selected. We follow the same overall principle by evaluating the disagreement between candidate model trajectories generated along the profile likelihood paths. However, the previous studies do not specify how the parameter space was discretised when constructing these trajectories. Here, we adopted a simple and reproducible discretisation scheme: we pick 10 evenly spaced values for each parameter across its plausible range. In each iteration, if a parameter is deemed practically identifiable, we only consider the points withing the estimated profile likelihood confidence interval. If no point falls within this interval, then we include the midpoint of the confidence interval to ensure coverage. For each selected value, we compute the corresponding constrained optimum using maximum likelihood estimation, generating at maximum a collection of 10 × *N* profile likelihood models across the *N* parameters.

To select the next experimental measurement location, the algorithm evaluates the variability among the generated model trajectories. Specifically, for each candidate location, it computes the variance of the predicted values. The candidate location with the highest variance is selected, as it is assumed to offer the greatest potential to improve parameter identifiability [12, 13]. In the original studies, this procedure was applied to single-output models, where the variance is a scalar quantity.

To adapt the benchmark algorithm to settings with multiple outputs, we replace the variance by the generalised variance which corresponds to the determinant of the covariance matrix. Furthermore, since the noise in our system depends on the magnitude of the observations, we also consider a scaled variant that accounts for the signal-to-noise ratio, by dividing the disagreement metric by the product of the estimated noise variances.

The estimation of the noise variance for truncated normal distribution is detailed in Appendix A. We obtain the relative variance by dividing the predicted output generalised variance by the estimated noise variance at each location, ensuring that the variances are of the same scale. Using two-dimensional outputs as an example, let **y** = [**y**_1_, **y**_2_]^*T*^ represent the matrix of predicted model outputs for the sampled parameters, and *σ*_1_ and *σ*_2_ denote the corresponding estimated noise variances. The mathematical expression for our relative variance **R** is then defined as follows:

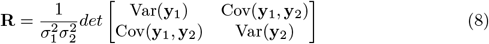

This metric allow us to jointly consider two outputs with different magnitudes of noise when making a data collection decision. Throughout the rest of the paper, we refer to the benchmark algorithm which has generalised variance being replaced by this relative-variance criterion as the *Scaled Benchmark*.

## 4 Empirical results

In this section, we demonstrate the advantages of E-ALPIPE over the benchmark algorithm and Random Sampling based on examples with one and two outputs. We first introduce the mathematical models used in this paper, define the noise distributions, and explain how we customise them for our research. The key difference between the examined methods lies in the data collection process: E-ALPIPE and the benchmark algorithm actively select data, whereas the random sampling method collects random settings from the candidate pool for each experiment. The three performance metrics used here are as follows:

1. **Success rate and speed of determining practical identifiability:** For the same number of collected data points, we compare the percentage of replications that achieve practically identifiability. For each replication, we record the iteration at which the model first becomes reliably practically identifiable, defined as the second of two consecutive iterations that both satisfy the identifiability criterion. The reported value is the mean of these iteration numbers across all replications.
2. **Width of the profile likelihood confidence interval:** We compare the profile likelihood confidence interval widths after a fixed number of 20 iterations.
3. **Point estimate quality:** We evaluate the quality of the MLE point estimates for the parameters after 20 iterations, by integrating over the absolute differences between the predicted function’s response and the true function response. The specifics of this calculation are detailed later in Section 4.4.

In practice, if a new data point provides information that conflicts significantly with the existing dataset, this can introduce new uncertainty about the parameter values. This could happen, e.g., if the new observation is an outlier. To enhance the reliability of our results, we consider an iteration to be identifiable only if it is identifiable across two consecutive iterations. This means that we require at least one extra data point to confirm our decision on practical identifiability.

In Experiment 1 below, all the results are averaged across **100** replications, while the number of replications is **30** in Experiments 2 and 3 due to the higher computational complexity of those two experiments. We terminate a run either when we reach stable practical identifiability or we have collected 50 data points (the default maximum for evaluating all algorithms). Each experiment commences with one observation at the same location which can be defined by the user. In this paper, the first observation for Experiment 1 is at *t* = 3, and *t* = 10 hours for Experiments 2 and 3. Then, iteratively additional data points are collected using each method. The practical identifiability and profile likelihood assessments are generated only from the second data point onwards. Finally, to assess statistical significance, we use the Wilcoxon rank sum test, which does not assume normality and is therefore more appropriate for our output distributions.

### 4.1 Models used in Experiments

We use three example models for the experiments in this paper. The first model is a sum of exponentials, which was used in [12]. The other two models are based on a nonlinear bioreactor, which was adapted from the microbial growth model described in [26]. We consider two versions of this bioreactor model, one with a single output, and one with two outputs.

#### 4.1.1 Sum of two exponentials

The first model considers the sum of two exponentially decaying components, with decay rates *β*_1_, *β*_2_ in units of hr^−1^. Only their combined total is observed at discrete time points, while both components are assumed to start at 1 when *t* = 0. The system can be formulated as a set of ODEs (Eq. 9) which have an analytical solution (Eq. 10), as previously described in [12]:

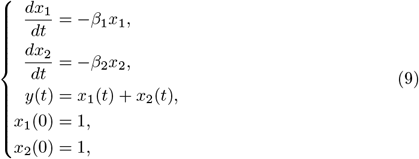

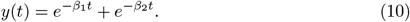

Since the model output *y*(*t*) is symmetric with respect to *β*_1_ and *β*_2_, interchanging the two parameters leads to identical observations. To ensure practical identifiability of individual parameters and to avoid obtaining two equivalent MLEs, we define their admissible ranges to be non-overlapping as follows:

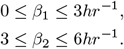

For our experiment, the true parameters are defined as *β*_1_ = 2 hr^−1^ and *β*_2_ = 4 hr^−1^. Observations are generated from the true model plus noise. The noise distribution is chosen to be a truncated normal distribution with mean 0 and standard deviation *σ* = 0.005, which ensures the observed values are non-negative.

The candidate times for measurement are from *t* = 1 to *t* = 10 hours in time steps of 0.1 hours. This experiment will be referred to as Experiment 1 in the following.

#### 4.1.2 Microbial growth model

While the sum of two exponentials model is simple and its ODEs are linear in parameters, we also present examples based on a nonlinear bioreactor model. This model was described by [26], with the following mathematical form:

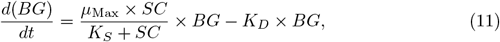

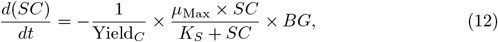

where *µ*_*Max*_ is the maximum growth rate, *K*_*S*_ refers to the 50% saturation constant for the substrate concentration, *K*_*D*_ refers to the decay rate and Yield_*C*_ is the yield coefficient. Note that the time dependence “*t*” is dropped from the states for the sake of notational brevity. The initial conditions used are as follows: *BG*(0) = 1*g/L* and *SC*(0) = 30*mM*. The model is structurally identifiable based on the Taylor series approach for SIA given that the initial conditions are known [27].

To account for realistic biological and operational variability, we define wide parameter bounds containing the true values as follows:

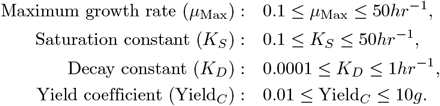

To evaluate performance under different conditions, we consider the following two setups:

- **Experiment 2** has a single output where only *bacterial growth (BG)* is observed. The only unknown parameters are *µ*_Max_, *K*_*D*_, and Yield_*C*_, while *K*_*S*_ is fixed at 30 mM (see below for details).
- **Experiment 3** has two outputs where both *bacterial growth (BG)* and *substrate concentration (SC)* are observed. In this case, all model parameters are treated as unknown.

The true function that simulates bacterial growth and substrate concentrations is based on parameter values adapted from [26]. All of the algorithms applied are tasked with establishing practical identifiability for the model parameters. The parameters in each experiment and their true values are given as follows:

- **Experiment 2:** *µ*_Max_ = 1 hr^−1^, *K*_*D*_ = 0.1 hr^−1^, and Yield_*C*_ = 0.5 g
- **Experiment 3:** *µ*_Max_ = 0.5 hr^−1^, *K*_*S*_ = 30 hr^−1^, *K*_*D*_ = 0.05 hr^−1^, and Yield_*C*_ = 0.6 g.

These parameter values fall within biologically plausible ranges and are chosen to ensure that no negative values are generated within the simulation time frame. The decision to fix *K*_*S*_ was based on a practical identifiability analysis conducted for Experiment 2 using the `MATLAB` tool `Data2Dynamics` [19], which determined that either *µ*_Max_ or *K*_*S*_ had to be fixed in order for Experiment 2 to be practically identifiable. The choice to fix *K*_*S*_ was further justified by its low sensitivity within the model (see Appendix B). The candidate times for measuring in both experiments are from *t* = 1 to *t* = 60 hours, with time steps of 0.5 hours. All time ranges were selected to capture the system’s full dynamic behaviour, while the step sizes ensure a sufficiently smooth resolution of the dynamics. These settings can be freely adjusted to suit specific experimental designs.

The standard deviation for the noise is based on the typical ranges of the observed output. For Experiment 2 which only has bacteria growth as the output, the standard deviation *σ*_*BG*_ of the noise distribution is set to 2. For Experiment 3, *σ*_*BG*_ is set to 0.5 and *σ*_*SC*_ to 1.6. A larger noise level is used in Experiment 2 to increase the difficulty of establishing practical identifiability, which allows us to better distinguish algorithm performance. The noise distribution is again chosen to be a truncated normal distribution.

A sensitivity analysis was performed to evaluate how model output variances respond to changes in parameters. Results for Experiments 1, 2, and 3 are presented in Appendix B.

### 4.2 Speed of determining practical identifiability

Based on the results shown in Figs. 1, 2 and 3, the E-ALPIPE algorithm exhibits a significantly faster increase in identifiability probability across iterations compared to the other two methods. Initially, E-ALPIPE achieves a much steeper ascent in all experiments, suggesting that it is more efficient in achieving practical identifiability results with fewer iterations. This indicates that our algorithm is more effective in selecting data points that provide significant information on model practical identifiability.

**Fig 1.**
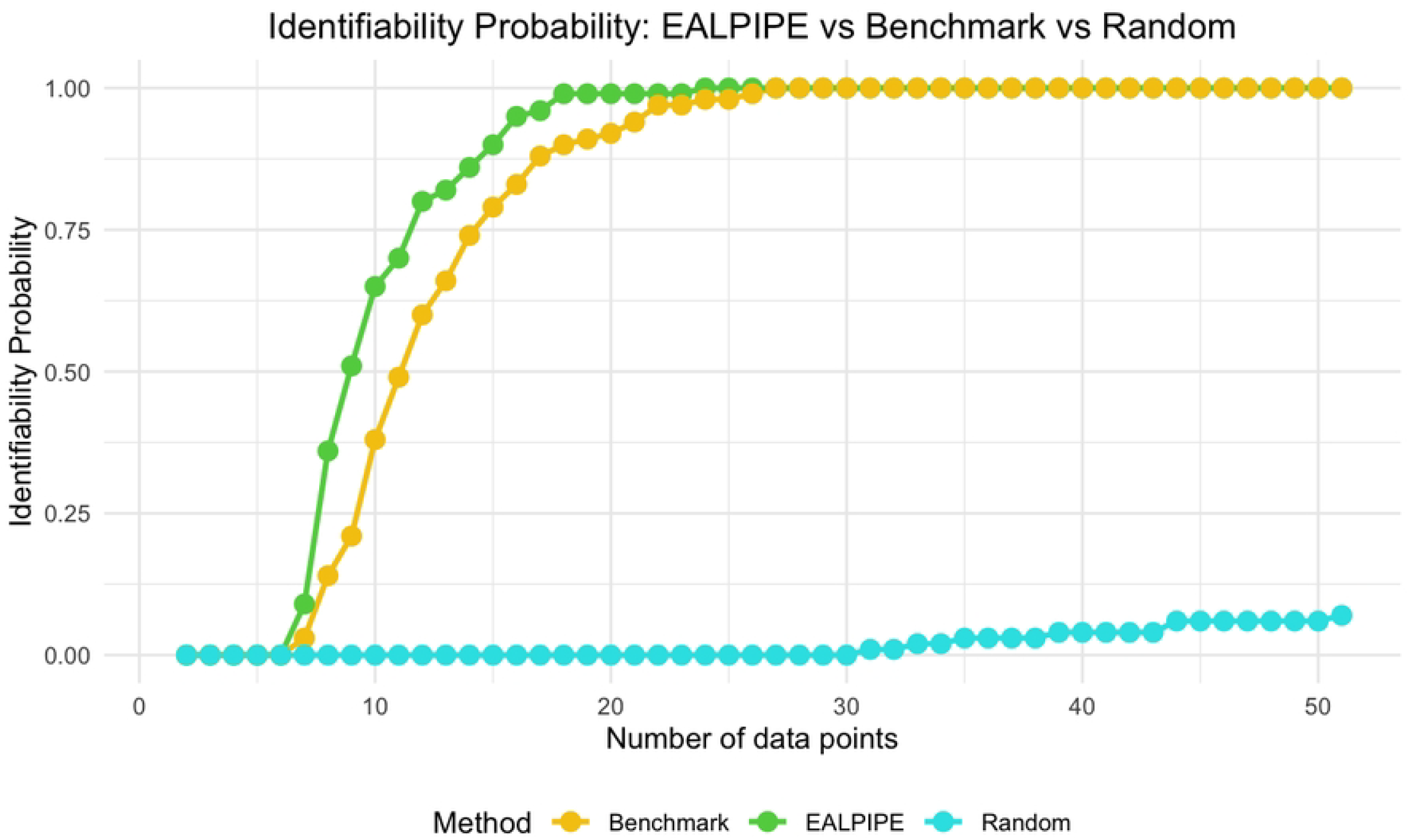
Identifiability probability for Experiment 1. Probability of achieving practical identifiability as a function of the number of collected data points for three algorithms: E-ALPIPE, Benchmark, and Random sampling. The number of data points includes the initial point, so the x-axis ranges from 1 to 51.

**Fig 2.**
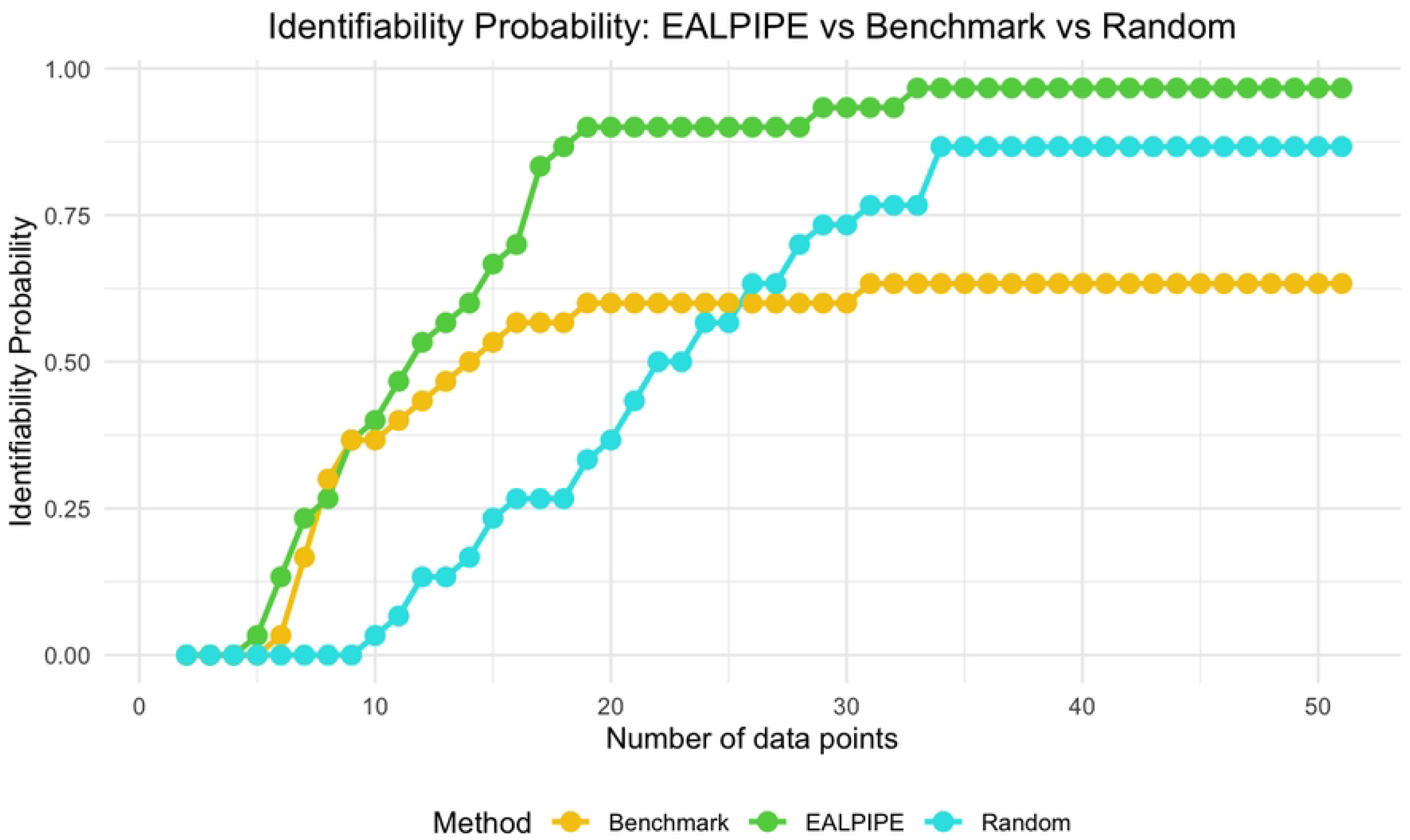
Identifiability probability for Experiment 2. Probability of achieving practical identifiability as a function of the number of data points collected for three algorithms: E-ALPIPE, Benchmark, and Random sampling. The x-axis includes the initial data point, so it ranges from 1 to 51.

**Fig 3.**
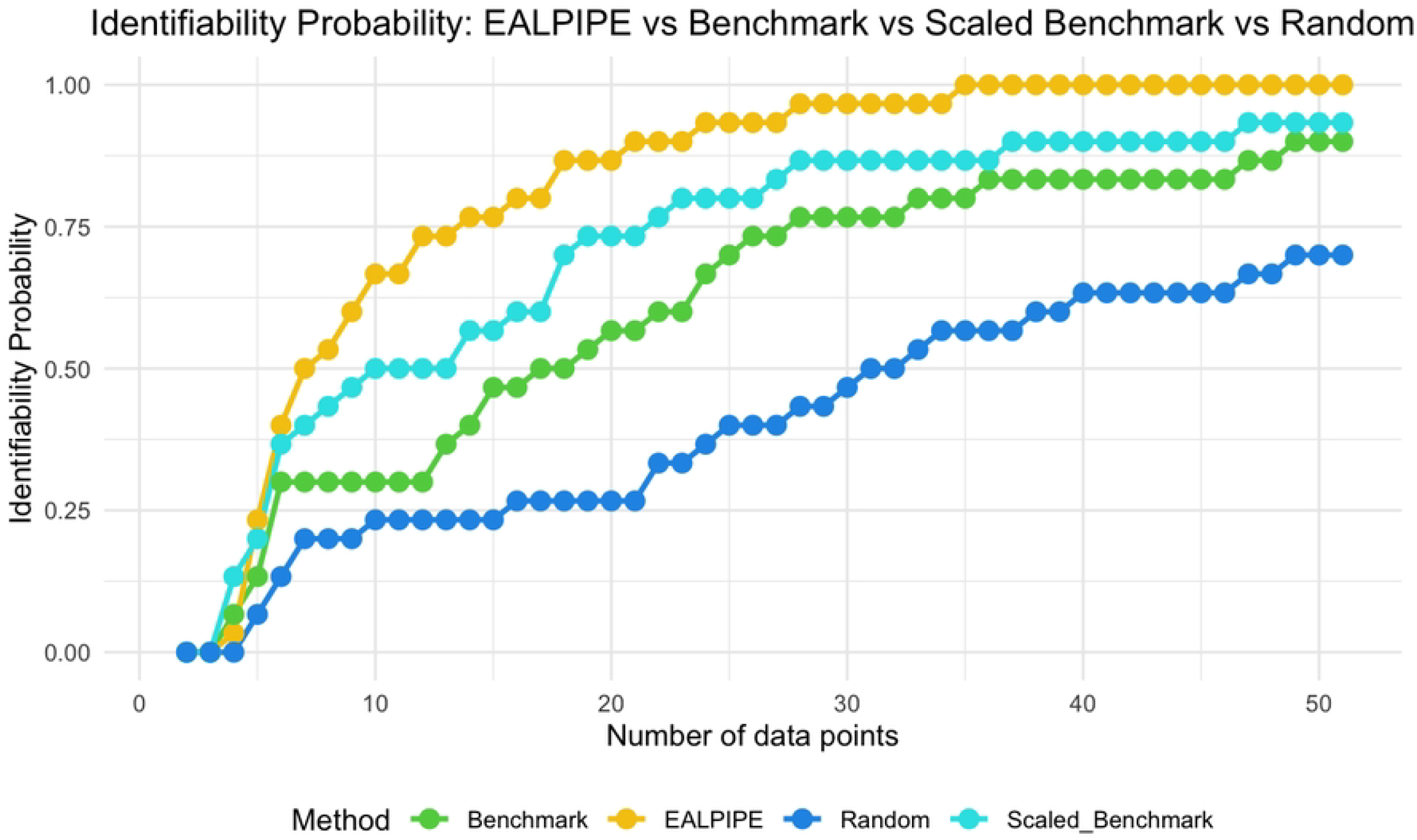
Identifiability probability for Experiment 3. Probability of achieving practical identifiability as a function of data-set size for four algorithms: E-ALPIPE, Benchmark, Scaled Benchmark, and Random sampling. The x-axis includes the initial point, so it ranges from 1 to 51.

We note that in Fig. 2, the benchmark algorithm performs worse than the random baseline. This unexpected result may be due to the benchmark repeatedly selecting locations with high variance, leading to oversampling of those regions. In contrast, while E-ALPIPE also focuses on informative areas, it demonstrates more exploratory behaviour. The selected sampling locations for each method are visualised as heatmaps in Appendix C.

In Fig. 3, the scaled benchmark outperforms the original benchmark. This highlights the benefit of normalising the disagreement criterion by the estimated noise variance, which helps prevent the selection from being biased toward outputs with inherently higher noise. Without this adjustment, the benchmark may be misled by outputs that exhibit large variability but provide limited information for identifiability.

We also report the average number of iterations required to establish practical identifiability in Table 1. For replications that failed to achieve practical identifiability, we assign as value the maximum number of iterations when computing the average. We highlight the method with the smallest average number of iterations required to achieve practical identifiability. An asterisk (*) is used to denote methods that are statistically significantly better than all others.

**Table 1.** Average number of iterations to achieve practical identifiability. The average number of iterations required to confirm practical identifiability is shown across three experiments over 4 methods.

| Method | Experiment 1 | Experiment 2 | Experiment 3 |
| --- | --- | --- | --- |
| E-ALPIPE | **9.46\*** | **13.2** | **9.97** |
| Benchmark | 11.7 | 24.8 | 20.4 |
| Scaled Benchmark | — | — | 15.4 |
| Random | 49.2 | 24.7 | 30.5 |

### 4.3 Width of the profile likelihood confidence interval

To evaluate each algorithm’s ability to produce precise parameter estimates, we assess the width of the profile likelihood confidence intervals (CIs) after each algorithm has collected exactly 20 observations. Note that, for a fair comparison across methods, we fix the number of observations to 20, so that each algorithm is evaluated using the same amount of information. In all experiments, we do not terminate the algorithm early even if practical identifiability is achieved before reaching 20 observations. The CI width serves as a proxy for estimator precision; narrower intervals suggest greater precision of the estimated parameter values. If a sampling strategy fails to achieve practical identifiability within 20 observations, we assign the full parameter search interval (as defined in Section 4.1) as its CI width for the purpose of averaging. Summary statistics of the resulting CI widths and their associated standard errors are presented in Tables 2, 3, and 4. In each column, the value with the smallest mean is highlighted. The results show that E-ALPIPE works best in most cases, although a Wilcoxon signed-rank test shows that most differences between the best and second best method are not significant, with the exception of E-ALPIPE and Random sampling, which yield a significantly narrower CI in Experiment 3 for parameter *K*_*S*_ and *K*_*D*_, respectively. The isolated case where Random sampling performs best is not sufficient to demonstrate its overall effectiveness, as it performs significantly worse than the other methods in most other settings. These results demonstrate that E-ALPIPE consistently achieves the narrowest profile likelihood intervals across most of the parameters and experiments, indicating greater estimator precision under the same data budget.

**Table 2.** Confidence interval width for Experiment 1. Mean width (± standard error) of the 95% confidence intervals for parameters *θ*_1_ and *θ*_2_ across different methods.

| Method | CI($\theta_1$) ($hr^{-1}$) | CI($\theta_2$) ($hr^{-1}$) |
| --- | --- | --- |
| E-ALPIPE | $0.114 \pm 2.88 \times 10^{-3}$ | **$1.13 \pm 3.33 \times 10^{-2}$** |
| Benchmark | **$0.109 \pm 3.68 \times 10^{-3}$** | $1.24 \pm 4.38 \times 10^{-2}$ |
| Random | $0.54 \pm 6.89 \times 10^{-2}$ | $3.00 \pm 0.00$ |

**Table 3.**
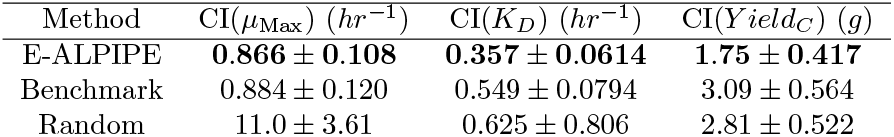
Confidence interval width for Experiment 2. Mean width (± standard error) of the 95% confidence intervals for parameters *µ*_Max_, *K*_*D*_, and *Y ield*_*C*_, across different sampling strategies.

| Method | CI($\mu_{Max}$) ($hr^{-1}$) | CI($K_D$) ($hr^{-1}$) | CI($Yield_C$) ($g$) |
| --- | --- | --- | --- |
| E-ALPIPE | **$0.866 \pm 0.108$** | **$0.357 \pm 0.0614$** | **$1.75 \pm 0.417$** |
| Benchmark | $0.884 \pm 0.120$ | $0.549 \pm 0.0794$ | $3.09 \pm 0.564$ |
| Random | $11.0 \pm 3.61$ | $0.625 \pm 0.806$ | $2.81 \pm 0.522$ |

**Table 4.** Confidence interval width for Experiment 3. Mean width (± standard error) of the 95% confidence intervals for parameters *µ*_Max_, *K*_*S*_, *K*_*D*_, and *Y ield*_*C*_ across four methods.

| Method | CI($\mu_{\text{Max}}$) ($hr^{-1}$) | CI($K_S$) ($hr^{-1}$) | CI($K_D$) ($hr^{-1}$) | CI($Yield_C$) (g) |
| --- | --- | --- | --- | --- |
| E-ALPIPE | **0.276 ± 0.0234** | **33.2\* ± 2.86** | 0.0153 ± 3.67$\times 10^{-4}$ | **0.0926 ± 9.61$\times 10^{-4}$** |
| Benchmark | 0.354 ± 0.0208 | 42.9 ± 2.84 | 0.0221 ± 8.70$\times 10^{-4}$ | 0.112 ± 3.54$\times 10^{-3}$ |
| Scaled Benchmark | 0.336 ± 0.0220 | 40.1 ± 2.73 | 0.0189 ± 4.34$\times 10^{-4}$ | 0.101 ± 2.20$\times 10^{-3}$ |
| Random | 0.408 ± 0.0251 | 51.5 ± 2.58 | **0.0121\* ± 3.87$\times 10^{-4}$** | 0.106 ± 2.96$\times 10^{-3}$ |

### 4.4 Assessing the point estimate quality

Another way to assess the quality of our sampling algorithms is to evaluate whether the data acquired by each sampling strategy yields parameter estimates that are capable of reproducing the true system dynamics. The predicted function value at each iteration is obtained by substituting the MLEs into the given model. To reduce the risk of convergence to local optima, we apply a multi-start optimisation approach when estimating the MLEs. In this section, we describe how we define prediction quality and report the corresponding experimental results. As in the previous section, each algorithm is terminated after exactly 20 observations have been collected to ensure a fair and consistent comparison across methods.

To compute the difference between the predicted and true functions over discrete time, we look at the output over a sequence of time points *T*, corresponding to the candidate time points specified in Section 4.1. At each time point *t* ∈ *T*, we calculate the absolute difference between the predicted value 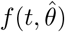 and the true value *f* (*t, θ*_true_), and then average these differences over all time points:

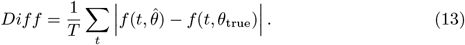

This metric represents the mean absolute deviation between the predicted and true values across all time points in the candidate set. Tables 5, 6, and 7 report the mean and standard error of this metric for each observable and sampling strategy across the three experiments, based on 20 collected observations per replication. The method which has the smallest average absolute difference is highlighted.

**Table 5.** Averaged absolute difference in predicted observable for Experiment 1. Mean absolute difference (± standard error) between predicted and true observable *y*(*t*) across three methods.

| Method | Diff<sub>y</sub> |
| --- | --- |
| E-ALPIPE | **1.58$\times 10^{-4}$ ± 9.63$\times 10^{-6}$** |
| Benchmark | 1.72$\times 10^{-4}$ ± 1.11$\times 10^{-5}$ |
| Random | 1.02$\times 10^{-3}$ ± 1.68$\times 10^{-4}$ |

**Table 6.** Averaged absolute difference in predicted output for Experiment 2. Mean absolute difference (± standard error) between predicted and true observable for BG across three sampling methods.

| Method | Diff<sub>BG</sub> |
| --- | --- |
| E-ALPIPE | **0.601 ± 0.0700** |
| Benchmark | 0.659 ± 0.0719 |
| Random | 1.21 ± 0.210 |

**Table 7.** Averaged absolute difference in predicted outputs for Experiment 3. Mean absolute difference (± standard error) between predicted and true outputs for BG and SC across four sampling methods.

| Method | Diff<sub>BG</sub> | Diff<sub>SC</sub> |
| --- | --- | --- |
| E-ALPIPE | **$0.179 \pm 0.078$** | $0.120 \pm 0.0540$ |
| Benchmark | $0.275 \pm 0.0363$ | **$0.105 \pm 0.0117$** |
| Scaled Benchmark | $0.254 \pm 0.0210$ | $0.113 \pm 0.0113$ |
| Random | $0.386 \pm 0.0530$ | $0.307 \pm 0.0420$ |

Although the Wilcoxon rank sum test indicates that the performance difference between the best and second best approach are often not statistically significant, E-ALPIPE consistently achieves the lowest or near-lowest average absolute differences in nearly all cases. This suggests that E-ALPIPE provides more accurate model dynamics under a fixed data budget in most practical scenarios.

## 5 Discussion

The experimental results presented demonstrate the effectiveness of the E-ALPIPE algorithm across three criteria relevant to practical identifiability analysis. In particular, E-ALPIPE outperforms the benchmark algorithm and random sampling method in terms of the speed with which practical identifiability is achieved, the precision of parameter estimates as measured by confidence interval widths, and the accuracy of system dynamics reproduction.

Across all three experiments, E-ALPIPE showed a faster increase in the probability of practical identifiability (probability of practical identifiability within 10 iterations is greater than 0.7) over the number of data points collected compared to Benchmark, Scaled Benchmark, and Random sampling (Figs. 1, 2, and 3). This indicates that the algorithm is more efficient at selecting informative data points that improve parameter practical identifiability. The improvement is particularly pronounced in Experiments 2 and 3, where E-ALPIPE required significantly fewer iterations on average to establish practical identifiability (Table 1). Specifically, in Experiment 2, the average number of iterations required by E-ALPIPE was less than 50% of that required by the other two methods. In Experiment 3, E-ALPIPE achieved practical identifiability using 42% fewer data points than the next best method, Scaled Benchmark.

In terms of estimator precision, in most experiments E-ALPIPE achieved narrower profile likelihood confidence intervals than the alternatives when the same amount of data were collected (Tables 2–4). This reflects the algorithm’s ability to guide data collection toward regions of the parameter space that reduce uncertainty.

Regarding point-estimate quality, which is quantified by the time-averaged absolute difference between true and predicted trajectories, E-ALPIPE again performs best in most cases (Tables 5–7).

To further understand how the algorithms prioritise sampling locations, we conducted a sensitivity analysis and present heat maps of the chosen observation against time (see Appendix B and C). The benchmark algorithm tends to focus on regions of high local sensitivity, potentially leading to over-sampling narrow informative regions. In contrast, E-ALPIPE demonstrates a more exploratory behaviour, distributing observations more broadly across the design space, which may contribute to its robustness and overall performance across different scenarios.

Since E-ALPIPE evaluates the profile likelihood only at the boundaries—rather than over the full parameter space—it also achieves substantial reductions in computational cost. This efficiency makes it especially suitable for computationally expensive models, where full-profile evaluations become prohibitive.

In summary, E-ALPIPE provides a resource-efficient, computationally feasible approach to experimental design through practical identifiability, consistently delivering faster identifiability, tighter confidence intervals, and accurate system reproduction under a fixed data budget.

## 6 Conclusion

Our results demonstrate that E-ALPIPE is an effective and resource-efficient strategy for active data collection in practical identifiability analysis. Across three synthetic case studies, the algorithm consistently achieved practical identifiability more quickly, produced narrower profile-likelihood confidence intervals, and reproduced system dynamics more accurately than a state-of-the-art benchmark and random sampling. These advantages arise from E-ALPIPE’s greedy information-gain criterion, which targets the practical identifiability by directly maximising the disagreement between the current predictive model and a constrained optimised model evaluated at the parameter boundaries. Our algorithm maintained its performance across models that differ in structure and dimensionality, underscoring its generalisability.

For experimentalists, these gains translate directly into reduced laboratory costs and time. Practical identifiability sets a lower bound on the data collection required for trustworthy parameter estimation; accelerating its confirmation therefore increases the robustness of model-based study design in bioengineering.

### Limitations

The present bisection search for profile-likelihood CIs assumes unimodal profiles and may result in wider than necessary confidence intervals for strongly multimodal posteriors. Also, all evaluations to date have been carried out on synthetic data; validation with real biological measurements is still required.

### Future work

Future work could extend E-ALPIPE to (i) multimodal profile likelihoods, (ii) continuous-time experimental design, and (iii) time-series observations rather than single time point measurements. We also plan to integrate the method into ongoing microbial bioprocess studies, providing an empirical real-world test of its cost-saving potential.

In summary, E-ALPIPE combines computational tractability with strong statistical performance, offering a practical path toward data-efficient experimental design in computational biology.

## Acknowledgement

The first author gratefully acknowledges support by the Engineering and Physical Sciences Research Council through the Mathematics of Systems II Centre for Doctoral Training at the University of Warwick (reference EP/S022244/1).

## Data availability statement

All code used for implementing the algorithm, conducting experiments, and performing data analysis is available on a GitHub repository at https://github.com/lulu0120/E-ALPIPE-Algorithm. We have also used Zenodo to assign a DOI to the repository: 10.5281/zenodo.16411995

## Appendix

### A Calculation of noise variance under truncated normal assumption

To account for the physical constraint that measurements cannot be negative, we model the measurement noise using a normal distribution truncated below at zero. At each location *x*, we assume the measurement *Y* follows a truncated normal distribution, *Y* ~ 𝒩 (*µ, σ*^2^) while restricting *Y* ≥ 0. Letting *ℓ* = 0 be the lower limit and 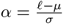, the density is proportional to the original normal PDF for values *Y* ≥ *ℓ*. Suppose *µ* is the predicted value from the MLE at that location and *σ* is a given standard deviation, then the estimated variance of *Y* is given by

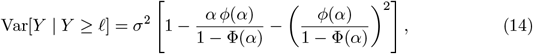

Where 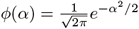, and 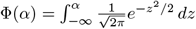 denote the standard normal probability density and cumulative distribution functions, respectively.

We estimate the noise variance at each sampling location as follows:

1. Evaluate the model prediction 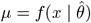, where 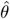 is the current MLE.
2. Compute *α* = (*ℓ* − *µ*)*/σ*.
3. Evaluate *ϕ*(*α*) and Φ(*α*).
4. Substitute into Eq. (14) to compute the variance of the truncated normal distribution.

This noise variance estimate is used to normalise model uncertainty in both the E-ALPIPE and Scaled Benchmark algorithms, where the estimated noise level informs the sampling selection stage.

### B Sensitivity analysis

A sensitivity analysis was performed in R based on the Sobol Indices approach [28]. In short, the total variance of a model’s output is separated and attributed to each of the model parameters and inputs of the model system. Sobol Indices provide a quantitative way of analysing the sensitivity of the model outputs based on the model parameters, where higher values indicate that the variance of the model’s outputs are sensitive to these parameters at a particular location, and low values indicate that the variance of the model’s output is less sensitive to these specific parameters.

For Experiment 1 (Fig. 4A) the Sobol indices for both *θ*_1_ and *θ*_2_ are above 0.9 throughout the entire experiment duration. For the bacteria growth observations in Experiments 2 and 3: For the BG output (Fig.4 B and C), the parameter *K*_*D*_ has the highest Sobol indices which are above 0.9. The parameter *Y ield*_*C*_, has Sobol indices above 0.5 during the first five hours of the experiment duration but then this decreases to less than 0.3 afterwards. The parameter *K*_*S*_ has the lowest Sobol indicies (*<* 0.2) overall for both the bacterial growth observation and also substrate concentration (Fig.4 D). The parameter *µ*_MAX_ has a higher Sobol index for the substrate concentration (*>* 0.9) output compared to the bacteria growth output.

**Fig 4.**
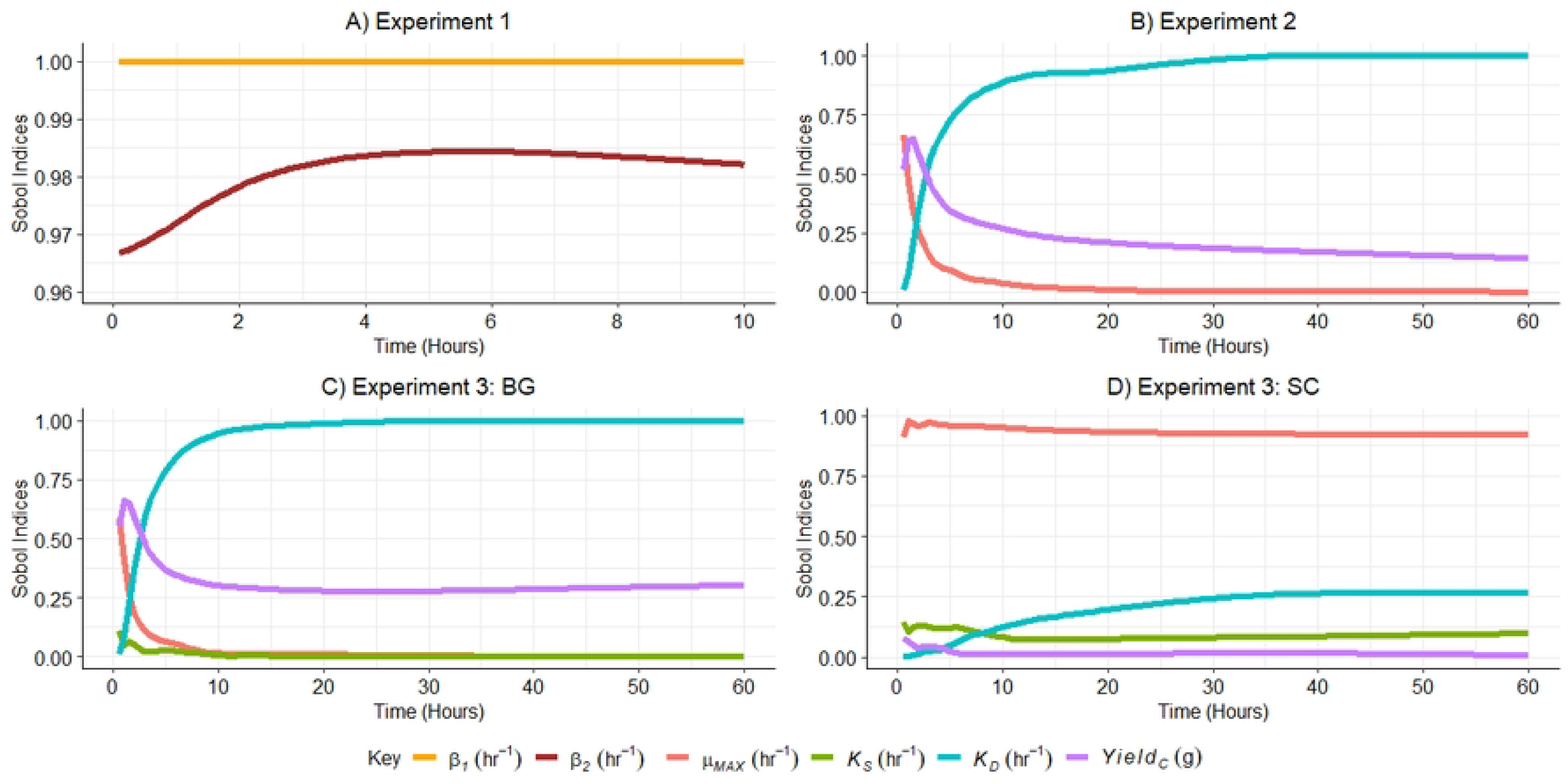
Sensitivity analyses for three experiments. Sensitivity analyses showing Sobol indices for Experiment 1 (A), Experiment 2 (B), and Experiment 3 (C and D). These results are based on model equations, experimental setups, and parameter bounds for each experiment (see Section 4.1).

### C Choice of sampling locations

To illustrate the measurement location selection across different sampling methods, Figures 5, 6, and 7 present heatmaps of selected locations aggregated over all independent replications, prior to achieving practical identifiability for all parameters. The plots demonstrate that in Experiments 2 and 3, E-ALPIPE, Benchmark, and Scaled Benchmark tend to select locations corresponding to regions of high Sobol sensitivity indices (Figure 4), confirming that these algorithms prioritise informative regions in the design space.

**Fig 5.**
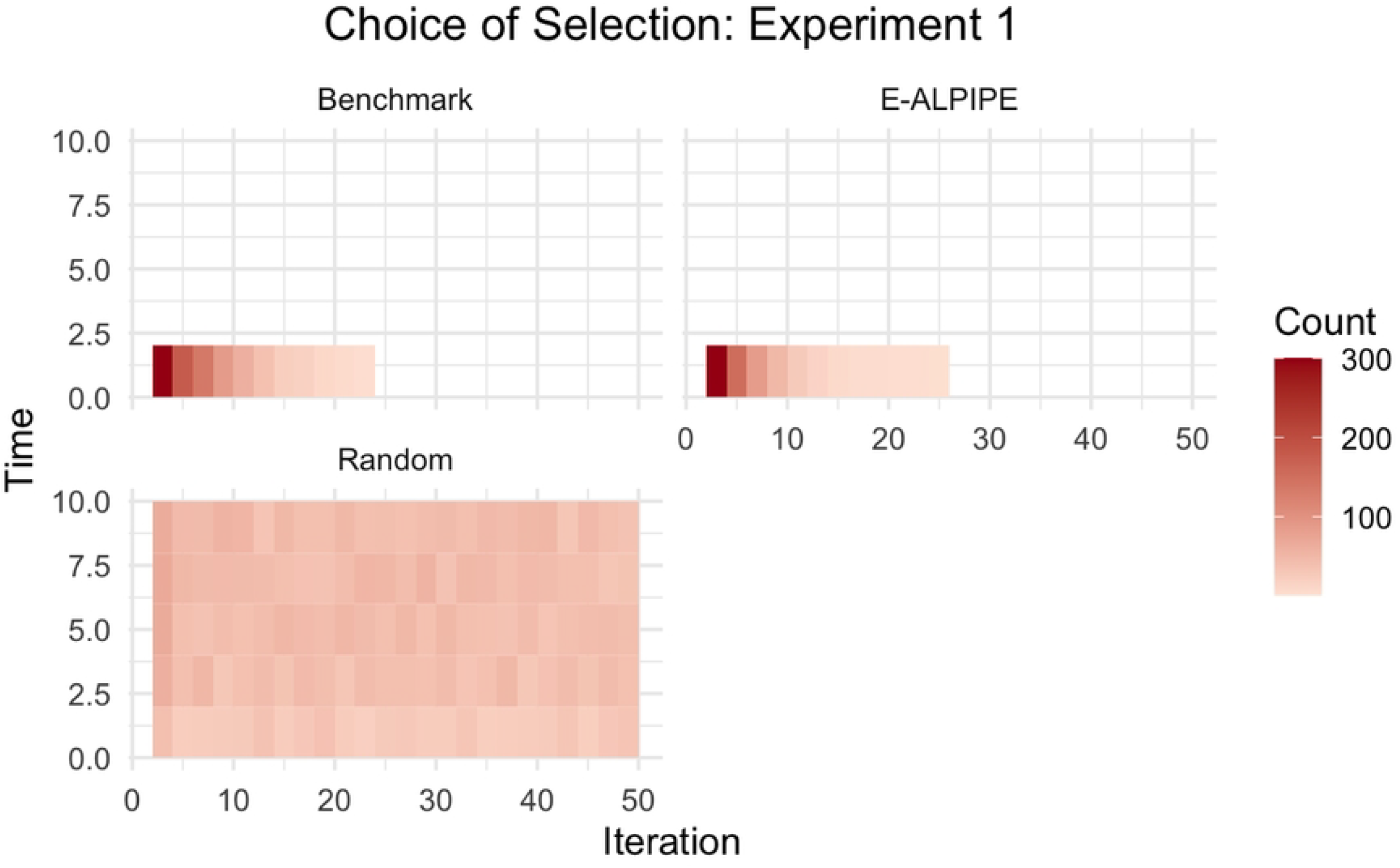
Sampling locations in Experiment 1. Heatmap of sampling locations selected by each algorithm in Experiment 1.

**Fig 6.**
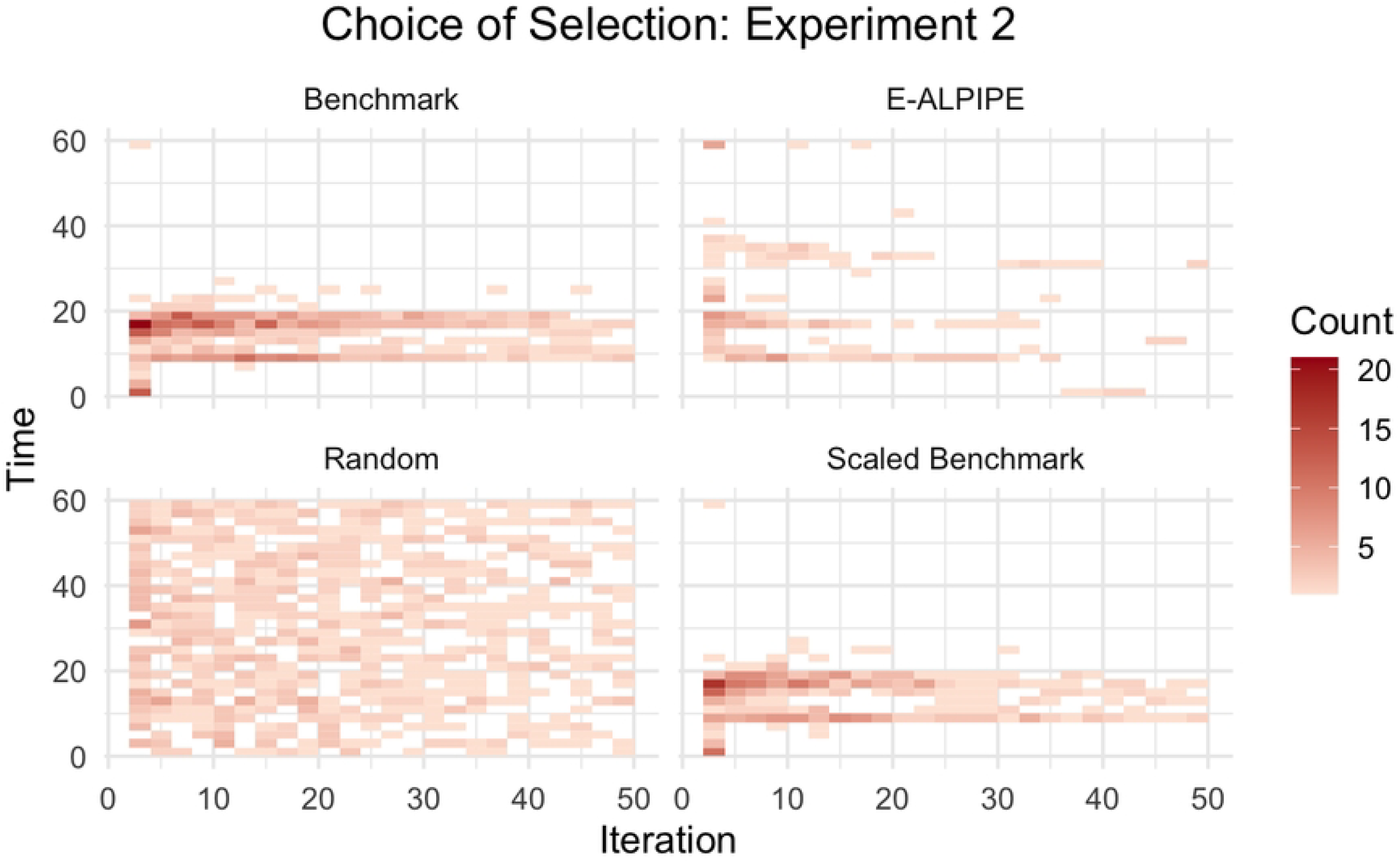
Sampling locations in Experiment 2. Heatmap of sampling locations selected by each algorithm in Experiment 2.

**Fig 7.**
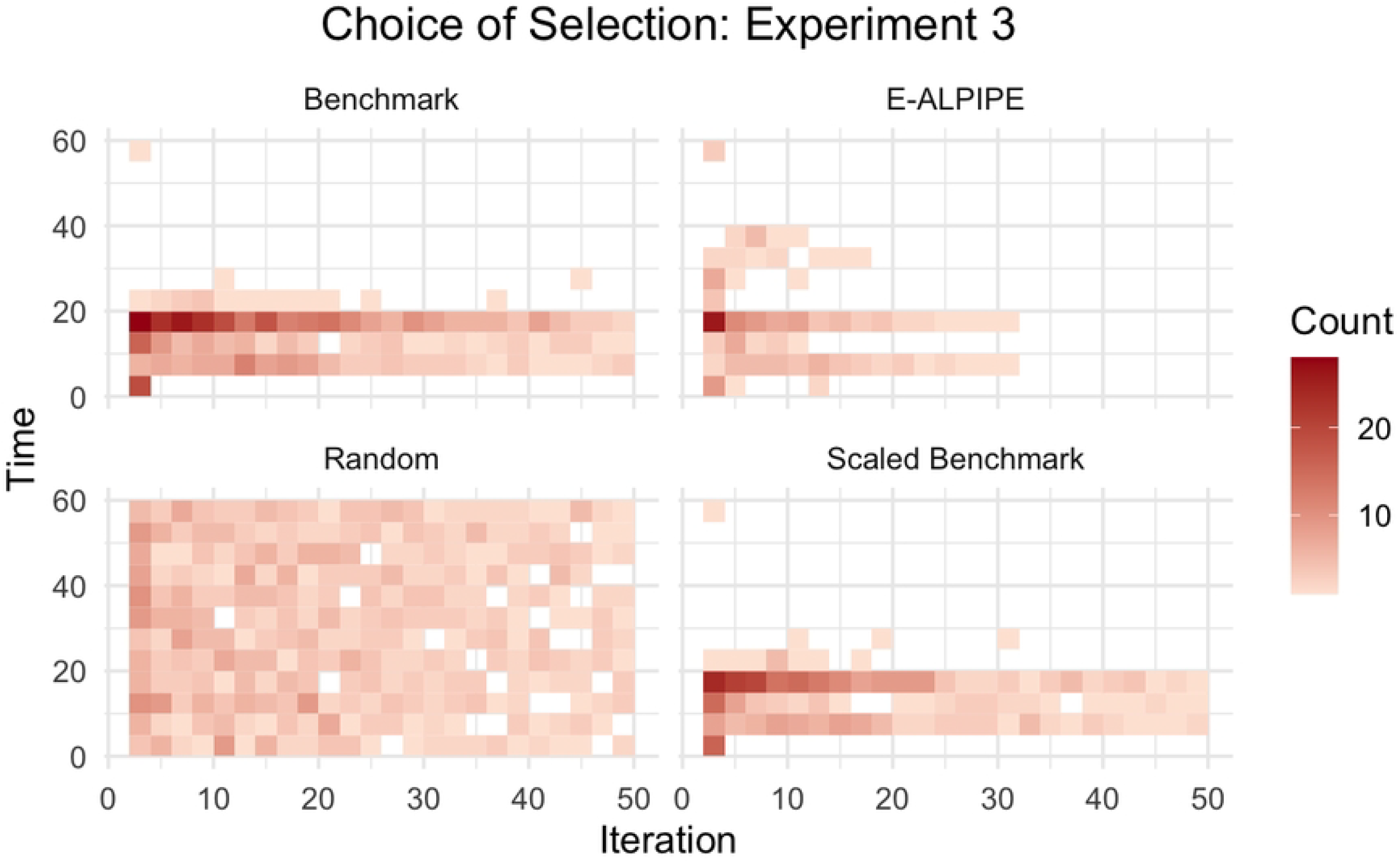
Sampling locations in Experiment 3. Heatmap of sampling locations selected by each algorithm in Experiment 3.

For Experiment 1, however, the sensitivity analysis suggests that regions beyond *t* = 4 hours provide limited information for *θ*_2_ due to the dominance of noise relative to the variability introduced by *θ*_2_. Both the Benchmark and E-ALPIPE algorithms correctly focus on earlier time points (before *t* = 2.5), effectively accounting for the noise level in their selection. Additionally, in Figure 6, the Benchmark algorithm exhibits more frequent repeated selections at certain locations, indicating potential over-exploitation of specific regions, whereas E-ALPIPE explores a broader set of informative regions. Note that E-ALPIPE and the Benchmark algorithm often achieve practical identifiability before iteration 50, hence the heatmaps are lighter in later iterations or stop entirely.

